# Injectable osteogenic microtissues containing mesenchymal stromal cells conformally fill and repair critical-size defects

**DOI:** 10.1101/362772

**Authors:** Ramkumar T. Annamalai, Xiaowei Hong, Nicholas Schott, Gopinath Tiruchinapally Benjamin Levi, Jan P. Stegemann

## Abstract

Repair of complex fractures with bone loss requires a potent, space-filling intervention to promote regeneration of bone. We present a minimally-invasive strategy combining mesenchymal stromal cells (MSC) with a chitosan-collagen matrix to form modular microtissues designed for delivery through a needle to conformally fill cavital defects. Implantation of microtissues into a calvarial defect in the mouse showed that osteogenically pre-differentiated MSC resulted in complete bridging of the cavity, while undifferentiated MSC produced mineralized tissue only in apposition to native bone. Decreasing the implant volume reduced bone regeneration, while increasing the MSC concentration also attenuated bone formation, suggesting that the cell-matrix ratio is important in achieving a robust response. Conformal filling of the defect with microtissues in a carrier gel resulted in complete healing. Taken together, these results show that modular microtissues can be used to augment the differentiated function of MSC and provide an extracellular environment that potentiates bone repair.

## Introduction

Bone has a remarkable capacity to regenerate through carefully orchestrated, cell-mediated repair processes [1]. However, healing in large and complex fractures is often impaired, leading to incomplete or functionally inferior bone regeneration. In some wounds, loss of the native vasculature and infection of the wound bed can further impair bone regeneration, resulting in a variety of pathologies, including delayed-, mal-, and non-unions. In such cases, therapeutic intervention to stimulate and accelerate the healing response is required. Bone grafting is a standard approach to this problem but needs invasive surgery to harvest and deliver the graft. Autologous grafts and flaps are limited in supply and can cause donor-site morbidity, including infection, hematoma, and pain [2]. Moreover, they are not suitable in 10-30% of cases due to difficulty in conforming the graft to the shape of the defect [3]. Allogeneic decellularized grafts and synthetic ceramic substitutes can also be used, but are biologically inferior compared to viable bone grafts due to the lack of cellular components. Although processes such as irradiation and lyophilization can reduce the risk of disease transmission from an allogeneic graft, they eliminate cellular components resulting in reduced osteoinductivity [4] and revascularization, resulting in higher bone resorption [5].

The ideal bone substitute would exhibit osteoconductive, osteoinductive, and osteogenic properties and would promote concomitant neovascularization of larger defects [4]. Materials-based approaches have been developed to promote osteoconductivity [1], and the immobilization and release of growth factors can be used as an osteoinductive cue. However, only cells can produce bone, and osteogenesis, therefore, requires either recruitment of endogenous cells or delivery of exogenous cells capable of forming bone. In large and ischemic defects, endogenous cell recruitment is impaired, and therefore cell transplantation may be required. Approaches in which appropriate progenitor cells are delivered in osteoconductive and osteoinductive microenvironments have particular promise because they combine the elements needed to regenerate bone even in challenging situations. Adult mesenchymal stromal cells (MSC) have been widely studied in this application because of their demonstrated osteogenic potential, but only limited success has been achieved in translation to the clinic [6]. Therefore, guiding the progenitor cell phenotype after transplantation is an intense field of research with the potential to create more robust methods to treat orthopedic bone defects.

Cell- and material-based strategies to bone regeneration have been successfully applied in the clinic [7], yet most approaches involve pre-formed scaffolds that require invasive surgery for implantation. Minimally-invasive delivery of cells and materials is preferable to minimize surgical complications and the possibility of injection [8]. In particular, delivery of moldable materials via injection has the advantage that they can conformally fill even irregularly-shaped defects. A variety of approaches have been developed to inject cells suspended in a protein or polymer solution, which can then be triggered to transform into a cell-laden hydrogel *in situ* [9]. Such hydrogels typically have relatively weak mechanical properties [10] unless they are chemically cross-linked, which can introduce cytotoxicity [8]. An alternate approach is to fabricate small tissue modules *ex vivo* by combining appropriate cells and extracellular matrix materials. This strategy has the advantage that the embedded cells can attach and remodel the matrix, and can be induced to proliferate and differentiate prior to being implanted via injection.

In this study, we combined adipose-derived MSC with a biomaterial matrix designed to mimic the native bone matrix and to provide osteoconductive cues. Native bone has a hierarchical structure composed of the protein collagen Type I, proteoglycans and glycosaminoglycans, and the calcium phosphate mineral hydroxyapatite [11]. We, therefore, developed a matrix that combined collagen (COL), which possesses motifs for cell adhesion that also guide cell function [12], with chitosan (CHI), a natural biopolymer that structurally and compositionally resembles GAG [11] and also improves mechanical properties and osteoconductivity [13, 14]. Hydroxyapatite (HA) was included to mimic the mineral phase of bone and to provide further biochemical and physical cues [15, 16]. This COL-CHI-HA composite material was used to create modular, spheroidal microtissues (60-100 μm in diameter) containing embedded MSC and designed to be delivered via injection through a standard needle. We characterized the morphology and composition of the microtissues, as well as the viability and phenotype of the embedded MSC. Microtissues were then implanted into a critical size cranial defect to investigate the effects of cell concentration, microtissue preparation volume, and localization of microtissues in the defect site on the quantity and quality of regenerated bone. This study demonstrates that modular microtissues can be an effective, minimally-invasive, cell-based approach to treating large bone defects.

## 2. Materials and Methods

### 2.1 Cells and Biopolymers

Adipose mesenchymal stromal cells (MSC) were harvested by digesting inguinal fat pads from 6-8 week old transgenic C57BL/6 ROSA^mT/mG^ mice (The Jackson Labs, Bar Harbor, ME) that express cell membrane-localized tdTomato (mT) fluorescence expression in all cells/tissues. The digestion solution consisted of 0.1 wt% collagenase in calcium- and magnesium-free phosphate buffered saline (PBS; Invitrogen). The harvested single cell suspension was filtered through a 70 μm cell strainer (Corning), suspended in Dulbecco’s Modified Eagle’s Medium (DMEM, Invitrogen) supplemented with 10% qualified FBS and 1% penicillin and streptomycin (Invitrogen) and cultured in tissue culture dishes. After 2-3 days of culture, adherent MSC were detached using TrypLE Express reagent (Invitrogen) and culture-expanded until passage 6 for use in experiments. Standard osteogenic supplements (OST) were added to the respective expansion media for differentiating MSC.

The biopolymers used to make osteogenic microtissues were chitosan (Protosan UP B 90/500, Novamatrix, Philadelphia, PA) and type I collagen (MP Biomedicals Inc., Santa Ana, CA). Chitosan stock solution was made through a two-step process. First 0.25 g of chitosan flakes were suspended in 25 ml of water and autoclaved at 121°C for 30 min. Then 30 μL of Glacial Acetic Acid (17.4 M) was added to the cooled suspension under sterile condition. The resulting solution (1 wt% chitosan in 0.02 N acetic acid) was stirred for seven days at 4 °C. The solution was then centrifuged at 10000 g to remove undissolved particles and used for making microtissues. Collagen stock solution was prepared by dissolving 250 mg of lyophilized collagen in 62.5 ml of sterile-filtered 0.02 N Acetic Acid. The resulting solution (0.4 wt% collagen in 0.02 N acetic acid) was stirred at 100-200 rpm for 7 days at 4 °C. The fully dissolved collagen solution was stored at 4 °C until microtissue fabrication.

### 2.2 Microtissue Fabrication and suspension-cultures

Chitosan-Collagen (CHI-COL) microtissues were fabricated using a modification of a water-in-oil emulsification method described previously [17, 18] (shown schematically in **Fig. 1A**). Preliminary matrix formulation experiments showed that a composite of 0.2 % collagen and 0.25 % chitosan exhibited minimal contraction (**Suppl. Fig. 1A**) while promoting early osteogenesis (**Suppl. Fig. 1B**). To prepare microtissues, cells were suspended in a mixture of solubilized chitosan, collagen, β-glycerophosphate (58 wt% in water, Sigma, St.Louis, MO), and glyoxal (68.9 mM in water, Sigma). All stock solutions were kept on ice prior to making the microtissue. For each 5 ml of the hydrogel mixture, 2.5 ml of the collagen stock (0.4 wt% in 0.02 N acetic acid), 1.25 ml of chitosan stock (1 wt% chitosan in 0.02 N acetic acid), 0.6 ml of β-glycerophosphate (58 wt% in water) and 60 μL of glyoxal (68.9 mM in water) was added and mixed thoroughly. Finally, 0.6 ml of cell suspension was added to the neutralized mixture and mixed thoroughly by gentle vortexing. To enhance the osteogenic potential of the microtissues, HA nanoparticles (0.1 to 2 wt%) were be added (**Supp. Fig. 1C**). Before adding to the mixture, the HA stock solution (300 mg/mL of PBS) was deflocculated by sonication. The cell-hydrogel mix was then dispensed dropwise into a stirred polydimethylsiloxane (PDMS, Clearco Products, Willow Grove, PA) bath kept on ice. The mixture was stirred by a dual radial-blade impeller at 800 rpm for 5 minutes to disperse the hydrogel composite into a fine emulsion in the PDMS. After 5 min, the temperature of the bath was increased to 37 °C and stirred for an additional 30 minutes to achieve full gelation of the microtissue droplets. The emulsion was centrifuged at 200 g for 5 min to separate the microtissues from the PDMS phase. The PDMS supernatant was removed without disturbing the pellet, which was then washed twice with complete culture media and collected by centrifugation at 150 g for 5 min. The microtissues were equilibrated in 5 ml of complete culture media to remove unbound β-glycerophosphate. The microtissue were then culture in vented tissue culture tubes in suspension culture inside a standard CO_2_ incubator at 37 °C.

**Figure 1.**
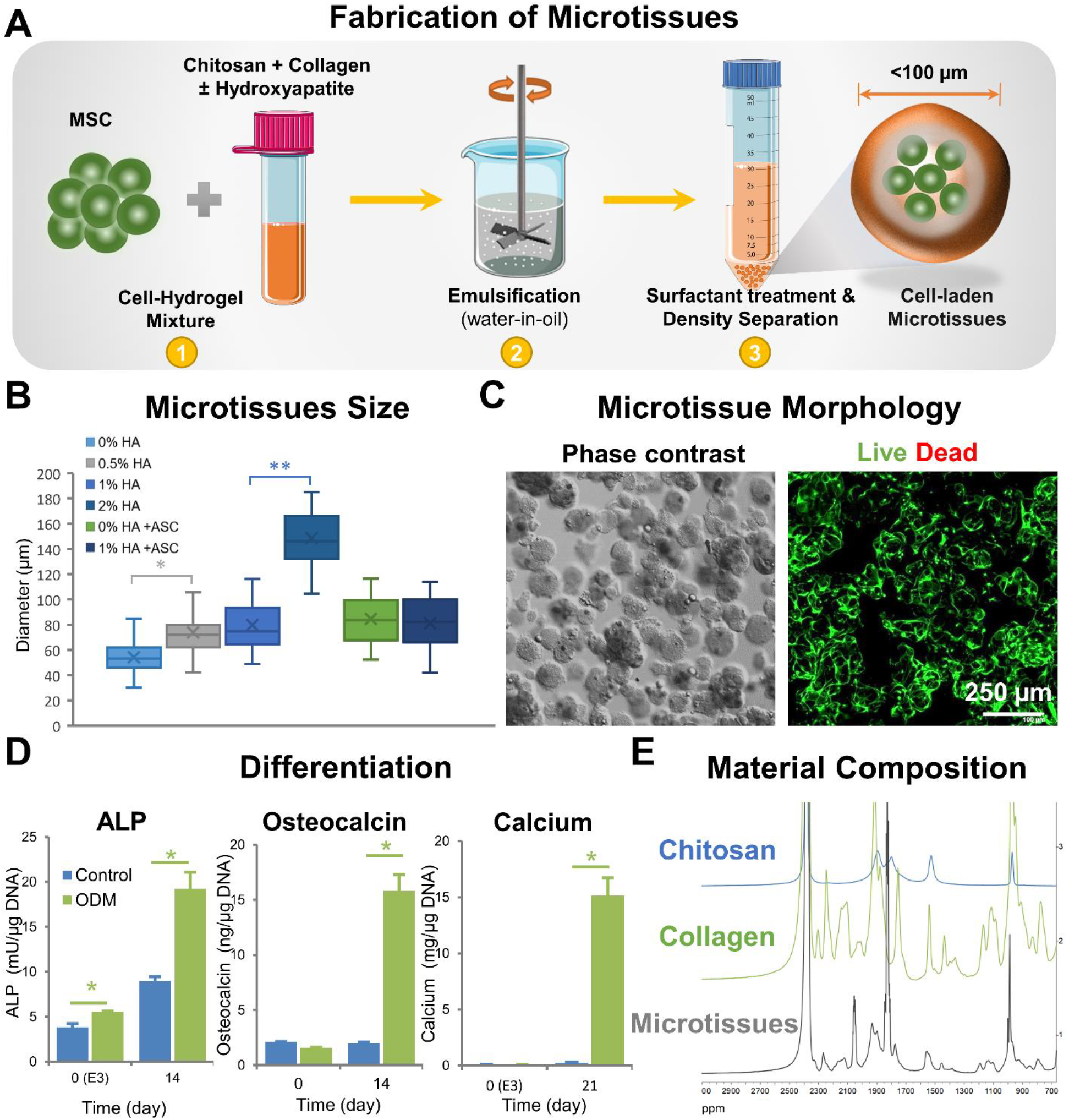
Fabrication and characterization of chitosan-collagen microtissues. A) Schematic of the water-in-oil emulsification process used to embed MSC within CHI-COL composite microtissues. B) Size distribution of CHI-COL microtissues as a function of HA content and cell loading. C) Phase contrast and fluorescence images showing the morphology of microtissues and vital staining (green) of embedded MSC. D) Expression of osteogenic markers by microtissues after osteogenic stimulation in vitro. E) ^1^H-NMR spectra of the microtissue matrix materials, demonstrating the presence of collagen and chitosan.

The morphology, sphericity, and size distribution of the microtissues were characterized using image stacks generated using a confocal microscope (Nikon Instruments, Melville NY). 3D projections of these stacks were then created using Imaris^®^ software (Bitplane, Belfast UK), and ImageJ software (National Institutes of Health) was used to characterize microtissue features.

### 2.3 Mouse calvarial defect and implantation

All animal procedures were performed in compliance with the Guide for the Care and Use of Laboratory Animals at the University of Michigan and approved by the Chancellor’s Animal Research Committee. All live animal surgical procedures were performed under 3-5% isoflurane/oxygen anesthesia. Immediately after induction, buprenorphine (0.05-0.1 μg/g body weight) and Xylazine (1.5 mg/kg) were administered for analgesia before the first surgical incision. To prevent corneal dryness, a bland ophthalmic ointment was applied during surgery. Sterile pharmacy grade saline was also administered (0.02 mL/g body weight) subcutaneously, to prevent dehydration. The calvarial defect was done as previously described [19]. Briefly, after cleaning the surgical site with Betadine, a 1.5-2 cm incision was made just off the sagittal midline in order to expose the right parietal bone. The periosteum enveloping the skull (pericranium) was removed using a sterile cotton swab. Using a diamond-coated trephine bit and copious saline irrigation, unilateral 4 mm full-thickness critical-sized calvarial defect was created in the non-suture associated right parietal bone, taking care to avoid dural injury. After injecting the microtissue construct into the defect, the skin was sutured with 6-0 vicryl suture, and the animals were monitored as per the established post-operative animal care protocols. Animals were kept under warm pads during recovery and observed for 6 hours before being returned to the animal housing facility. Animals were monitored twice daily for three days and weekly thereafter to ensure postoperative recovery. Buprenorphine was administered every 12 hours for two days postoperatively. Preliminary studies showed that microtissue formulations lacking an HA component failed to produce appreciable ossification in the calvarial defect (**Supp. Fig. 1**). Therefore, HA was included in all tested formulations.

### 2.4 In vivo imaging and analysis

In vivo bone formation was assessed with longitudinal micro-computed tomography (micro-CT) scans for 12 weeks. Micro-CT was performed using a high-resolution SkyScan 1176 small animal imaging system (Bruker, Billerica, MA) for up to 15 weeks. Images were reconstructed and bone volume formation was analyzed using a MicroView software (GE Healthcare, London ON, Canada). Quantification was performed by selecting for new, calcified bone only within the initial defect with a Hounsfield radiodensity of 1250 or higher based on Misch bone density classification [20]. Every mouse was scanned with a CT-phantom which included hydroxyapatite, water, and air for calibration.

To track the red fluorescent tdTomato (mT) from exogenously implanted MSC, an IVIS^®^ Spectrum in vivo imaging system (PerkinElmer, Shelton CT) and confocal microscope (Nikon) were used. At week 12 the animals were sacrificed by CO_2_ asphyxiation and cervical dislocation and imaged immediately after removing the skin to eliminate auto-fluorescence. The animal calvaria were placed in the imaging system and imaged for 2 seconds at small binning. The fluorescence at the calvarial injury site was quantified using Living Image 3.2 (PerkinElmer). The calvaria were then fixed in buffered alcoholic formalin solution (Z-Fix, Anatech) and stained for DAPI (Invitrogen). After rinsing twice in 10 mM PBS, samples were resuspended in fresh PBS and images were captured using a confocal microscope (Nikon).

### 2.5 NMR Analysis

Nuclear magnetic resonance (NMR) spectroscopy was employed to study the composition of the microtissue matrix as previously described [21]. Briefly, collagen-chitosan microtissue samples were collected, washed in PBS, dialyzed (8 kDa MWCO) against DI water for 48 hours at 4 °C and lyophilized to obtain dry samples. Then 2.0 mg of each sample was dissolved in deuterated water (D_2_O, 0.5 mL) containing 0.5% CD_3_COOD and dissolved at room temperature for 4 hours before analysis. ^1^H NMR spectra in D_2_O were recorded on 700 MHz Varian Mercury systems (Palo Alto, CA) at room temperature. ^1^H NMR spectrums were referenced using Me_4_Si (0 ppm), residual D_2_O at δ ^1^H-NMR 4.65 ppm. ^31^P-NMR was used to evaluate the phosphorous groups from β-GP.

### 2.6 Histology and imaging

After 12 weeks, animals were euthanized using CO_2_ asphyxiation and cervical dislocation to allow confirmation of microCT imaging results. The calvaria were harvested fixed, decalcified and paraffin embedded as described previously [22-24]. The embedded tissues were sectioned using a microtome into 6 μm thin sections and stained with Pentachrome, Hematoxylin and Eosin dyes for analyzing neo-vessel formation, mineralization, and bone morphology. For vascularization analysis, samples were immunostained for CD31 and evaluated for microvessels as described previously [25]. The sections were then mounted and imaged with a bright-field microscope (Nikon).

### 2.7 Ultrasound imaging-guided in-vivo injection of microtissue

To demonstrate minimally-invasive delivery, microtissues were injected into a premade defect using a 23 gauge needle guided by ultrasound imaging. Before the procedure, the animal was euthanized using CO_2_ asphyxiation to reduce pain and suffering. A 4-mm critical-sized calvarial defect was created, and the skin was immediately sutured as detailed above. The animal was placed on the examination stage and imaged with a high-resolution small animal ultrasound unit (Visualsonics, Toronto, Canada). Hair was removed from the skin above the calvarial defect, and acoustic gel was applied to couple the skin with the ultrasound probe. The microtissue were injected into the defect with a 23 gauge needle while the procedure was being recorded with the Vevo 708 scan head (55 MHz center frequency; 20-75 MHz -6 dB bandwidth; 30 μm axial resolution; 70 μm lateral resolution; 4.5 mm focal distance; 1.5 mm -6 dB focal depth; 100% transmit power) at a frame of 11 frames/s. 3D images of the defect before and after the injection were acquired by moving the ultrasound probe with a step size of 30 μm over an 8 mm range while scanning. Spectral ultrasound analysis was done to discriminate the native tissues and microtissue construct based on their mineral content as described previously [26, 27]. Briefly, grayscale values GS(y,z), acoustic scatterer diameter and acoustic concentration were extracted from the raw backscattered radio-frequency (RF) signals as described previously [26]. For power spectrum analysis, each of the RF scan line was segmented and a linear fit was applied to the calibrated gated power spectrum to determine the slope and mid-band fit (MBF). The effective scatterer diameter (a) was calculated from the slope, the geometry index, the center frequency of the imaging transducer, and bandwidth of the transducer.

### 2.8 Statistical analysis

All measurements were performed at least in triplicate. Data are plotted as means with error bars representing the standard deviation. Statistical comparisons were made using Student’s *t*-test and two-way ANOVA with a 95% confidence limit. Differences with p<0.05 were considered statistically significant.

## Results

### 3.1 Fabrication and characterization of chitosan-collagen osteogenic microtissues

The microtissue fabrication process and characterization of the resulting tissue modules is shown in **Figure 1**. The water-in-oil emulsification method (shown schematically in **Fig. 1A**) consistently produced spheroidal microtissues containing uniformly embedded MSC. Different mass ratios of chitosan-collagen (CHI-COL) hydrogels were tested to create stable microtissues through emulsification. The CHI-COL hydrogel formulation of 0.25 % CHI and 0.20 % COL yielded stable microtissues with minimal contraction (**Suppl. Fig. 2A**) while promoting early osteogenesis (**Suppl. Fig. 2B**). This selected CHI-COL formulation yielded microtissues with an average diameter of 54.3±12.1 μm (**Fig. 1B**). Addition of HA nanoparticles to enhance calcification increased the size of microtissues in a dose-dependent fashion, with the diameter of 0.5%, 1%, and 2% HA CHI-COL microtissues being 73.9±17.7 μm, 79.7±18.7 μm, and 148.6±21.4 μm respectively. Addition of 0.5 million cells/ml yielded microtissues with an average diameter of 84.6 ±18.2 μm. Increasing the cell concentration reduced the number of acellular beads, but the average diameter of the cell-laden microtissues remained constant. Microtissues with 0.5 million cells/ml and 1% HA had a diameter of 81.1±20.1 μm, and were not statistically significantly different from those containing only 0.5 million or only 1.0% HA.

In vitro characterization showed that the microtissues were stable and remained as individual modules in culture while maintaining high cell viability (**Fig. 1C**). Exposure to osteogenic supplements in culture resulted in clear increases in makers of osteogenesis over 14 days in culture, including the secretion of increased levels of alkaline phosphatase and osteocalcin, and the deposition of calcium mineral (**Fig. 1D**). NMR analysis of the matrix composition showed that collagen and chitosan have distinct spectra, and demonstrated the presence of both components in the microtissues (**Fig. 1E**). NMR analysis of the microtissues over time showed that the matrix composition was maintained over three weeks in culture (**Suppl. Fig. 3**).

### 3.2. Bone regeneration in critical-sized calvarial defect

Microtissue implants were tested for the ability to regenerate bone in a full-thickness critical size cranial defect in the mouse. In a first study, the effect of the cellular component was examined, as shown in **Figure 2**. Microtissues (created using 1 ml of initial hydrogel volume) with no cells, undifferentiated MSC, or osteogenically predifferentiated MSC (OD-MSC) were implanted for 15 weeks with regular periodic assessment by microCT and histological analysis at explant. Gross examination of the 3D reconstructed microCT images showed that by week 10, osteogenically predifferentiated microtissues had bridged >95% of the defect area (p<0.001, **Fig. 2A and Suppl. Fig. 4**). In contrast, microtissues with undifferentiated MSC showed significantly less bony bridging (p<0.05) with only ~50% coverage. Microtissues without cells showed the lowest degree of defect bridging, covering only ~30% of the defect area. Correspondingly, the total bone volume, bone mineral content, bone mineral density and bone volume fraction (bone volume/tissue volume) of the newly formed bone within the defect was significantly higher in the osteogenically predifferentiated microtissues compared to the acellular and undifferentiated MSC conditions (p<0.001, **Fig. 2B-E**). Histological analysis of the newly formed bone in osteogenically-predifferentiated samples revealed a lamellar and woven bone morphology, high collagen deposition (yellow staining) and structures resembling early marrow cavities (**Fig. 2F**). Microtissues containing undifferentiated MSC exhibited only minimal bone formation at the defect borders and generally low collagen deposition and were characterized by sparse bone spicules surrounded by fibrous connective tissue. The microtissues without cells had the lowest mineral content, though the defect was fully bridged with collagenous osteoid structures interspersed with blood vessels (**Fig. 2F**).

**Figure 2.**
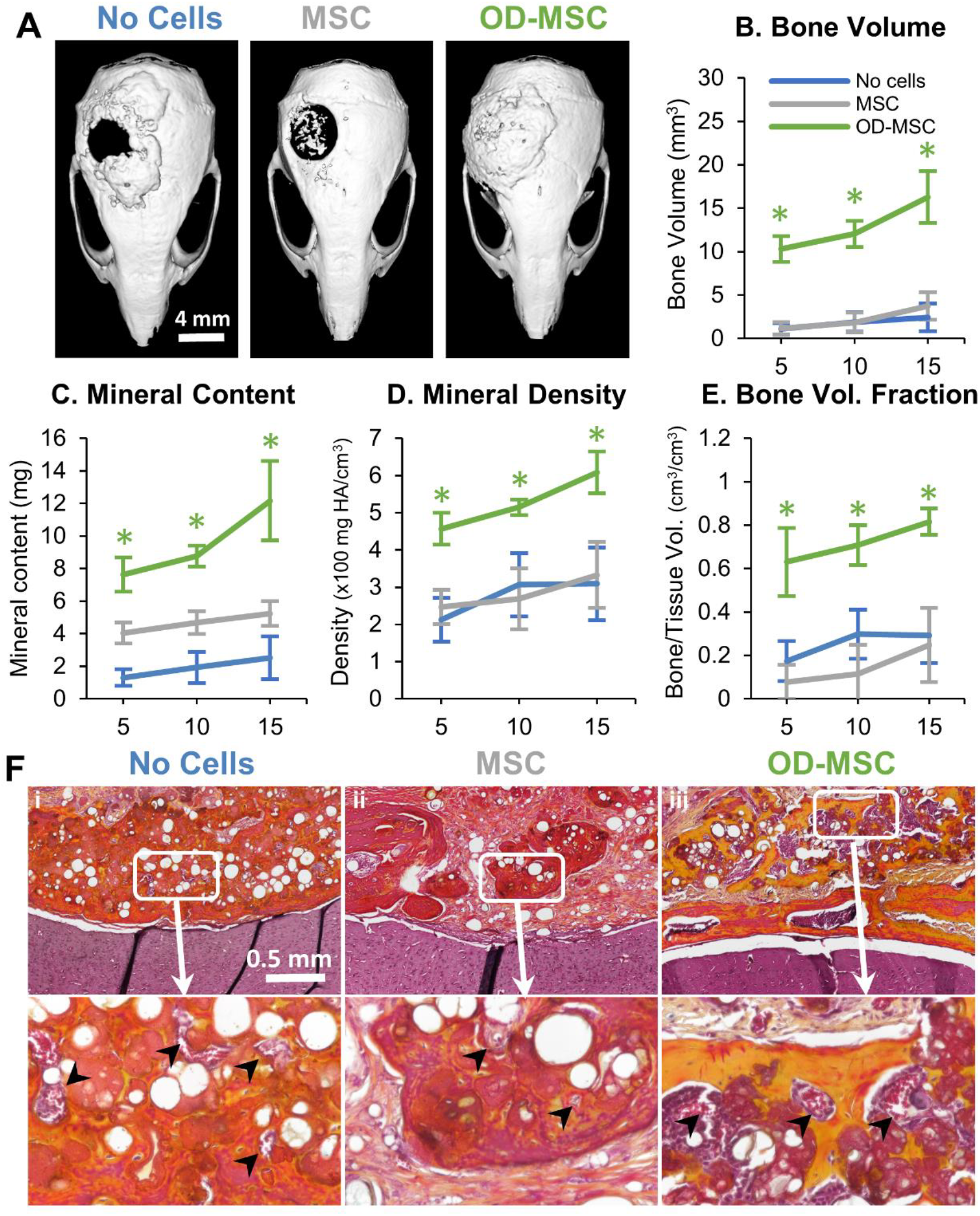
Bone regeneration in a critical-sized calvarial defect. A) Representative microCT images of bone formation in the defect region at 10 weeks. MicroCT data were analyzed to specifically assess new bone in the 4 mm defect site across implant replicates, and to obtain quantitative measures of: B) total bone volume, C) mineral content, D) mineral density, and E) bone volume fraction (bone volume/tissue volume). F) Histology images of newly formed bone in the defect site using Movat’s pentachrome staining. (Collagen fibers – yellow; fibrinoid and fibrin-bright red; nuclei – purple-black). The arrows indicate microvessels.

### 3.3. Influence of microtissue implant volume and cell concentration on bone formation

The robust and voluminous bone formation resulting from the implantation of osteogenically predifferentiated microtissues (OD-MSC) extended beyond the defect site and produced superfluous tissue adjacent to the defect. The volume of microtissues delivered was therefore varied to determine the most effective delivery bolus for regeneration of bone in the defect site. Implants were made using osteogenically predifferentiated microtissues created using 0.25 mL, 0.50 mL and 0.75 mL of initial hydrogel volume, at a constant cell concentration of 0.5x10^6^ cells/mL of the hydrogel. These microtissue preparations were implanted into a critical size cranial defect in the mouse, and the results are shown in **Figure 3**. Characterization of bone formation at 0, 3, 6, 9 and 12 weeks post-implantation using microCT showed that the volume of new bone decreased with decreasing initial volume of the microtissue implant (**Fig. 3A**). There was no statistical difference in bone volume between the highest and medium amounts tested, but new bone volume dropped significantly at the lowest implant volume (**Fig. 3B**). A similar dose-dependence was observed in bone mineral content (**Fig. 3C**), bone mineral density (**Fig. 3D**) and bone volume fraction (**Fig. 3E**), with lower implant volumes generally leading to less robust regeneration. Consistent with the radiographic analysis, histological analysis of the calvaria at 12 weeks showed less apparent formation of lamellar and woven bone as the microtissue implant volume decreased (**Fig. 3F**).

**Figure 3.**
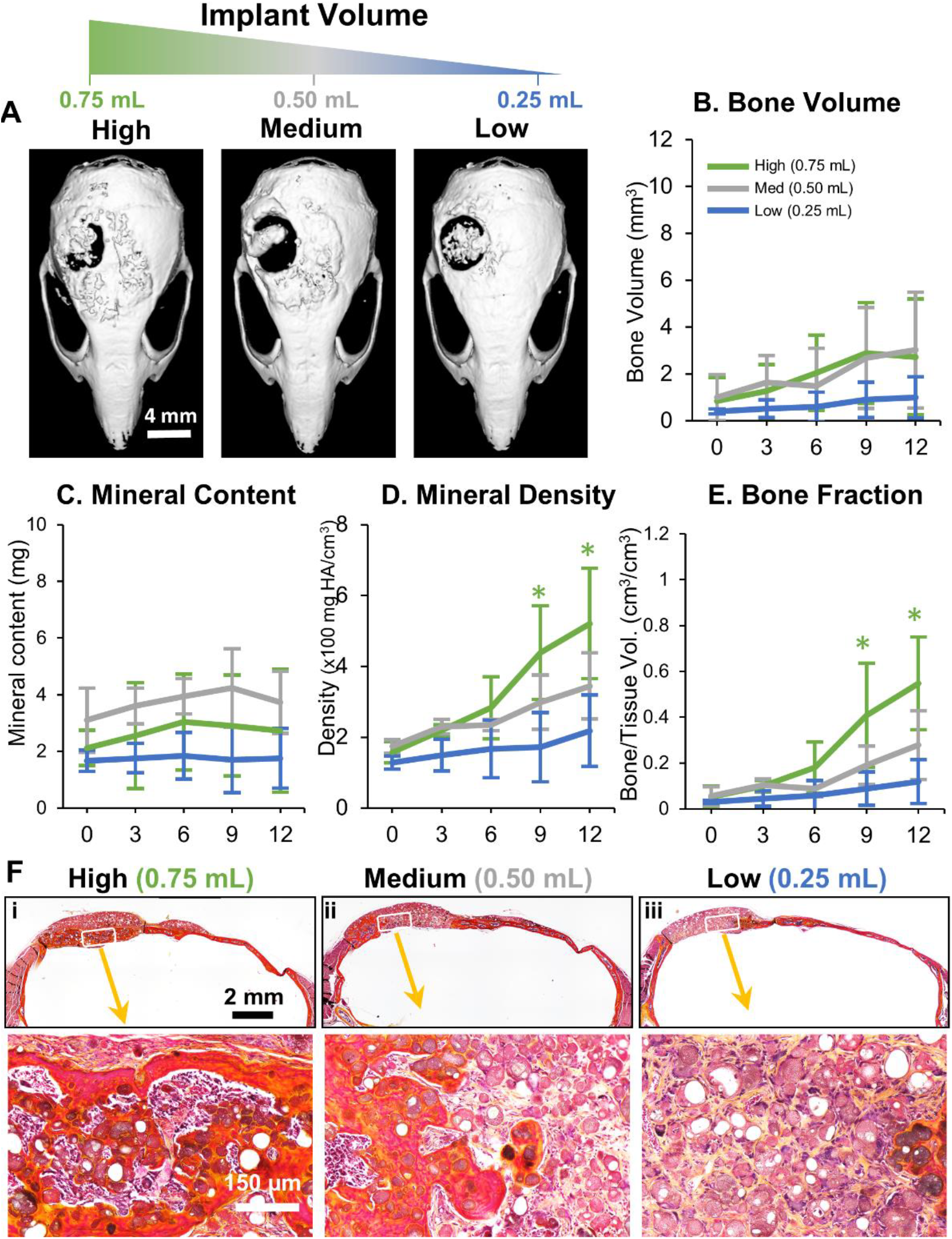
Influence of microtissue implant volume on bone formation. A) Representative microCT images of bone formation in the defect region at 12 weeks. MicroCT data were analyzed to specifically assess new bone within the 4 mm defect site across implant replicates, and to obtain quantitative measures of: B) total bone volume, C) mineral content, D) mineral density, and E) bone volume fraction (bone volume/tissue volume). F) Histology images of newly formed bone in the defect site using Movat’s pentachrome staining. (Collagen fibers – yellow; fibrinoid and fibrin-bright red; nuclei – purple-black).

The influence of the cell concentration used in the microtissue implants was also investigated, which keeping the implant volume constant. Microtissue implants containing OD-MSC microtissues made at concentrations of 1.0x10^6^, 2.0x10^6^ and 3.0x10^6^ cells/mL were studied in the mouse critical size cranial defect, as shown in **Figure 4**. Interestingly, an increase in cell concentration reduced the degree of ossification of microtissues and volume of bony structures in the defects (**Fig. 4A**) in a dose-dependent manner. Higher cell concentration resulted in significantly reduced new bone volume (**Fig. 4B**), bone mineral content (**Fig. 4C**), and total bone fraction (**Fig. 4E**), while the lowest cell concentration consistently outperformed the medium- and high concentration implants in these metrics. The bone mineral density, however, stayed essentially the same between the groups (**Fig. 4D**). Histological analysis revealed lamellar and woven bony structures with collagen deposition within the defect in all conditions, but larger areas of ossified regions were seen in implants at lower cell concentration (**Fig. 4F**).

**Figure 4.**
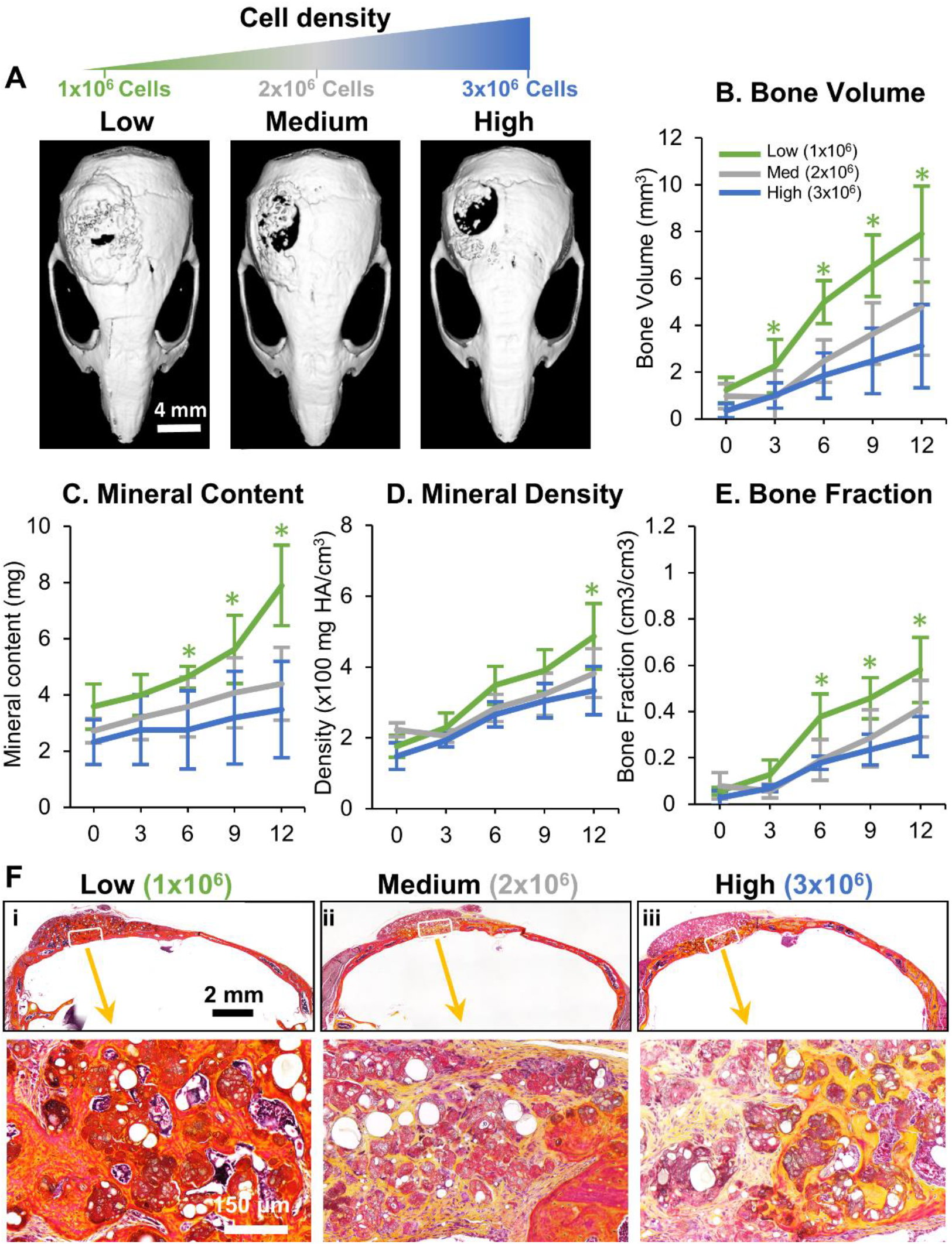
Influence of cell concentration on bone formation. A) Representative microCT images of bone formation in the defect region at 12 weeks. MicroCT data were analyzed to specifically assess new bone within the 4 mm defect site across implant replicates, and to obtain quantitative measures of: B) total bone volume, C) mineral content, D) mineral density, and E) bone volume fraction (bone volume/tissue volume). F) Histology images of newly formed bone in the defect site using Movat’s pentachrome staining. (Collagen fibers – yellow; fibrinoid and fibrin-bright red; nuclei – purple-black).

### 3.4. Delivery of microtissues in a fibrin carrier gel

The open nature of the critical size calvarial defect makes it difficult to contain microtissues only within the defect site. Therefore the effect of concentrating the microtissue population in the defect using a fibrin carrier gel was examined, as shown in **Figure 5**. Microtissues made with OD-MSC were suspended in a fibrin gel, and a plug of this material was prepared to fit the dimensions of the calvarial defect. These implants were compared to similar microtissues transplanted as a paste, and microtissues created with undifferentiated MSC served as an additional control. MicroCT imaging showed clearly that the carrier gel effectively contained the microtissues within the calvarial defect and conformed closely to the defect area (**Fig. 5A**). Within nine weeks the defect was bridged entirely in both the OD-MSC conditions with evidence of active remodeling. Quantitative analysis of microCT data showed no significant difference in the new bone volume between the OD-MSC microtissues implanted with or without a carrier gel and both groups significantly outperformed the implants with undifferentiated MSC (**Fig. 5B**). Likewise, the mineral density, mineral content and bone volume fraction were markedly higher in the OD-MSC implants, though there were no significant effects of the use of a carrier gel (**Fig. 5C-F**). Histological analysis revealed dense collagenous lamellar and woven bone formation in OD-MSC implants, compared to much looser and less defined structures in the undifferentiated microtissue implants (**Fig. 5F**). Analysis of vascularization of the newly formed tissues showed that implants with OD-MSC had significantly higher blood vessel area (**Fig. 5G**), with well-developed blood vessels perfusing the implant volume (**Fig. 5H**). Fluorescent imaging of the calvaria using an in vivo imaging system showed the presence of fluorescently tagged exogenous MSC in all conditions at week 12 (**Suppl. Fig. 5**), confirming the engraftment of the implanted cells.

**Figure 5.**
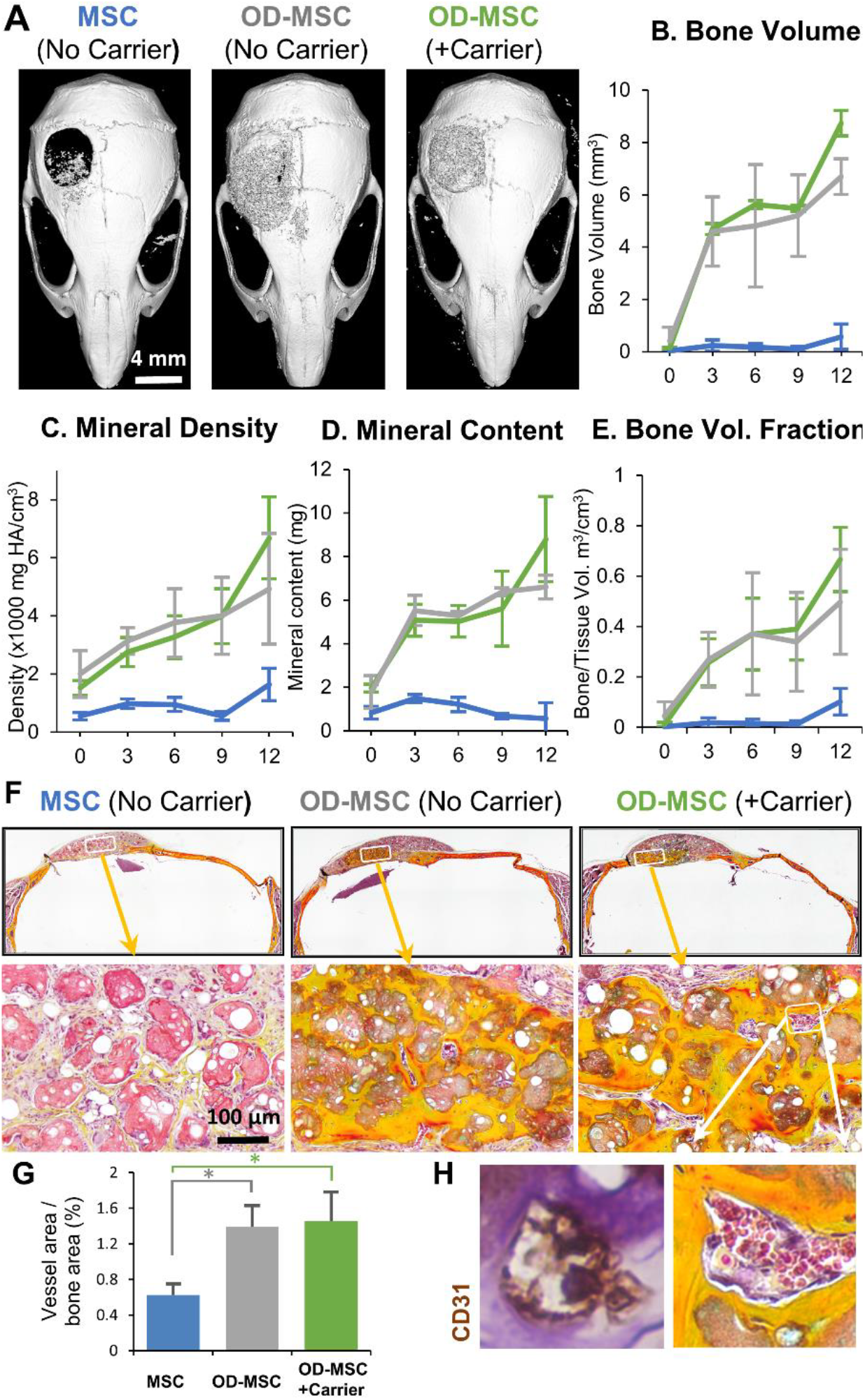
Effect of carrier gel to concentrate microtissue within the defect. A) Representative microCT images of bone formation in the defect region at 12 weeks. MicroCT data were analyzed to specifically assess new bone within the 4 mm defect site across implant replicates, and to obtain quantitative measures of B) total bone volume, C) mineral content, D) mineral density, and E) bone volume fraction (bone volume/tissue volume). F) Histology images of newly formed bone in the defect site using Movat’s pentachrome staining. (Collagen fibers – yellow; fibrinoid and fibrin-bright red; nuclei – purple-black). G) Blood vessel area with the defect was quantified from H) histology images, which showed well-developed vessels containing erythrocytes throughout the volume of OD-MSC microtissue implants.

### 3.5. Ultrasound-guided minimally invasive delivery of microtissues

The ability to deliver microtissue implants via injection was demonstrated by administering them directly through the skin to the calvarial defect in a mouse through a standard 23G hypodermic needle (330 μm inner diameter). Microtissue delivery was guided by high-resolution spectral ultrasound imaging to track progress in real time and to provide information on the composition of the implants (**Fig. 6A**). Ultrasound imaging enabled the creation of 3D spatial maps of the defect and the surrounding tissues, and periodic images showed the transfer of the microtissue paste into the implant side (**Fig. 6B, see Suppl. Info. for real-time video**). Moreover, the spectral ultrasound imaging (SUSI) technique applied in this study was also used to quantify the mineral content of the implanted tissues. Microtissues with varying HA concentration were imaged and analyzed, showing that the midband fit parameter varies linearly with mineral content (**Fig. 6C**). Similarly, analysis of the acoustic concentration in microtissue implants provides a spatial map of the mineral composition of the constructs (**Fig. 6D, Suppl. Fig. 6**). These non-destructive ultrasound imaging and analysis techniques can be used to monitor cell delivery, and potentially also can be applied to noninvasive and quantitative longitudinal monitoring of implant ossification and defect healing.

**Figure 6.**
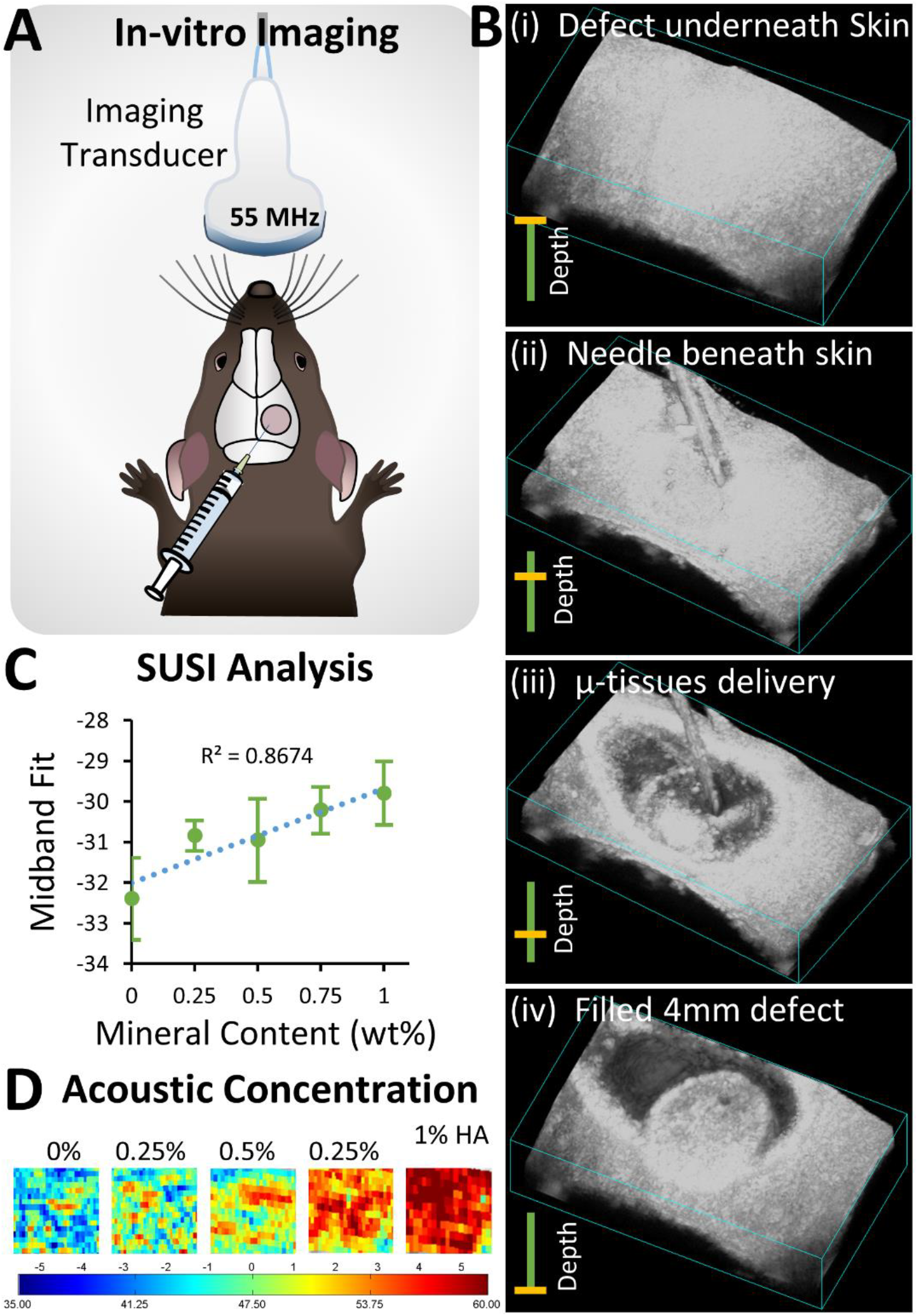
Ultrasound-guided, minimally invasive delivery of microtissues. A) Schematic of the monitoring microtissue implantation into the calvarial defect using high-resolution ultrasound imaging. B) 3D ultrasound image reconstructions showing injection of microtissues through the skin into the calvarial defect. C) Correlation of mineral content of microtissues and the midband fit parameter generated by spectral ultrasound imaging (SUSI). D) Heat maps of acoustic concentration generated by SUSI, showing spatial distribution and concentration of mineral in the microtissue implants.

## 4. Discussion

Despite progress in developing new treatments for complex fractures and large bone defects over the past decade, there remains an important need for stimulating bone regeneration in a variety of serious indications [28, 29]. There is a mismatch between the clinical demand for advanced therapies and the degree to which clinical translation has been achieved. Importantly, the desired properties of a tissue-engineered bone substitute depend on the intended clinical use. Fractures in large, load-bearing bones require mechanical stability and the regeneration of structurally strong bone. In other indications, the defect may not be load-bearing, but associated soft tissue loss and a challenging healing environment require a therapy that can potentiate the healing process. Therapies that can be delivered without major surgery are particularly attractive because they reduce the chance of infection and the burden of recovery. Injectable, cell-based approaches such as the one described here may therefore have particular value in treating large and complex bone defects.

The availability of an autologous cell source is an essential advantage for a cell therapy to avoid immune complications. The adipose-derived MSC used in this study can be easily obtained from the stromal vascular fraction of lipoaspirate [30]. These cells are highly proliferative [19, 31] and have verified potential to differentiate toward osteogenic, adipogenic, and chondrogenic lineages [32-35]. The extracellular matrix used in this study was chosen because of its structural and biochemical resemblance to the early bone matrix. It combines the protein collagen Type I with the GAG-like polysaccharide chitosan and further includes a hydroxyapatite mineral phase to nucleate bone formation. The process used to create cell-laden microtissues using these matrix materials is facile and allows tailoring of the size and composition of the microtissues to suit the intended application [36]. In particular, our studies demonstrated that microtissues made from MSC embedded in a CHI-COL-HA matrix maintain cell viability and are supportive of osteogenic differentiation in vitro. While the size of the microtissues depended to some degree on their composition, it was straightforward to create osteogenic microtissues with a diameter of 60-100 microns, which was the target range for modules to be easily injectable through standard needles.

The goal of this therapy was to fully bridge a large, critical size defect in the mouse calvarium. Interestingly, transplantation of acellular microtissues (i.e., the CHI-COL-HA matrix alone) for 15 weeks resulted in some bony tissue formation, but only in apposition to the native bone of the skull, while the defect was filled mainly with loose collagenous osteoid. These results suggest that the biomaterial alone is osteoconductive, but does not have strong osteoinductive properties. The incorporation of undifferentiated MSC into microtissues resulted in less collagen deposition, but created islands of calcified bone in the defect by week 15, and the total new bone volume within the defect was not significantly higher than that formed by acellular microtissues. In contrast, osteogenically predifferentiated MSC resulted in a very robust bone formation that completely bridged the defect, and in fact, extended beyond the boundaries of the defect when the implant was not constrained.

The difference in the effect of undifferentiated and predifferentiated MSC in microtissues after implantation may be a result of their distinct immunomodulatory and anti-inflammatory roles [37]. Adipose-derived MSC lack the HLA-DR receptor [38], an MHC class II receptor and antigen-presenter that is commonly implicated in graft loss. In addition, MSC are known to reduce inflammation through the secretion of prostaglandin E2 (PGE2), which can suppress lymphocyte action [39] and reprogram macrophages to an anti-inflammatory phenotype [40, 41]. However, it has been shown that pro-inflammatory signals secreted by macrophages are essential in the osteogenic function of MSC. For example, tumor necrosis factor-alpha (TNFα) and interleukin 6 (IL-6) are both critical pro-inflammatory cytokines that are secreted by activated macrophages, and both have been shown to play an essential role in the osteogenic commitment of adipose-MSC [42, 43]. Therefore, the anti-inflammatory effects of cytokines secreted by undifferentiated MSC may be responsible for the reduced bone regeneration observed when these cells are transplanted in microtissues.

In this study, OD-MSC were predifferentiated toward the osteogenic lineage using dexamethasone, a regulator of the osteogenic transcription factor Runx2 [44], and β- glycerophosphate, a phosphate source for mineralization. It has been shown that similarly predifferentiated MSC interact with macrophages after transplantation to improve defect healing [45]. In addition, priming of MSC toward the osteogenic lineage with potent biochemical factors such as bone morphogenetic protein-2 (BMP-2) [46], platelet-derived growth factor (PDGF) [47], and TNFα [48] prior to transplantation has been shown to be effective in regenerating bone. Taken together, these results suggest that there is a benefit to predifferentiation of MSC toward the osteogenic lineage prior to transplantation, not only to potentiate neotissue formation but also to promote synergistic interactions with the inflammatory cascade.

The effects of changing the implant volume and the cell concentration in implants were also investigated. Decreasing the volume of microtissues implanted had the effect of reducing lamellar bone formation, mineral deposition, and tissue remodeling. However, when the concentration of cells in the implant was increased, there was an apparent decrease in resulting bone formation and remodeling. Interestingly, both increasing the absolute implant volume and increasing the cell concentration per volume has the effect of transplanting a larger number of cells. However, only in the case where cells were transplanted with a correspondingly increased volume of extracellular matrix was there a positive effect on bone formation. It is possible that implants with a high cell density suffer from hypoxia [49-51], which can lead to tissue necrosis[52]. Alternately, there may be other beneficial effects of cell-matrix contacts that lead to better osteogenic outcomes. Overall, it appears that maintaining an appropriate ratio of cells to the matrix is essential in maximizing the regenerative potential of MSC.

Two modes of delivering microtissue implants were examined. A primary advantage of the modular microtissue format is that they can be transplanted minimally invasively through a standard needle. The injectability of microtissues was demonstrated under guidance and monitoring by high-resolution ultrasound imaging. Filling of the defect through the skin was facile. However, it was observed that implants of microtissue paste extended beyond the boundaries of the cranial defect and produced ectopic bone in the region. It should be noted that this issue is at least partially an artifact of the implant model used, since the geometry of the cranial defect it not conducive to retaining implanted material. Other indications, such as segmental defects surrounded by soft tissue, spinal fusions, and more enclosed cavital defects may be more appropriately filled with microtissue paste without loss of material. To address the issue of containment of microtissues in an open defect, a study was performed to examine the use of a polymerizable, fibrin-based carrier gel to entrap and restrain the microtissues. Fibrin was chosen as a carrier material because of its relative abundance, ease of handling, and wide use as a surgical sealant [53]. Also, fibrin has been shown to be effective in promoting angiogenesis [54, 55], as well as in creating vascularized tissues including bone [56, 57]. The fibrin gel carrier was effective in retaining the microtissues within the defect, and these implants resulted in similarly robust bone morphology, collagen deposition, and mineral content as unconstrained implants. In vivo imaging demonstrated that exogenous MSC were present at the implant site for the duration of the study.

This study demonstrates that injectable, MSC-laden chitosan-collagen microtissues can be used to bridge large bone defects with vascularized lamellar and woven bone. Osteogenic priming of the microtissues showed a clear benefit in accelerating bone regeneration. Importantly, the microtissue format allows osteogenic preculture and subsequent transplantation of microtissues without disrupting cell-matrix contacts. Therefore, the phenotype of cells in microtissues is likely to be preserved post-transplantation, and cells are more likely to engraft and survive in challenging healing environments. In contrast, transplantation of undifferentiated MSC in microtissues had an inhibitory effect on bone formation, possibly through mediation of the inflammatory response. The role of MSC phenotype and its interactions with the wound healing cascade are of great importance in developing advanced bone regeneration strategies, and there is an opportunity to capitalize on this knowledge. The chitosan-collagen biomaterial used in this study was osteoconductive, and studies on transplanted cell number indicated that the ratio of cells to the matrix in the implant is essential in achieving robust regeneration. Taken together, this work shows that microtissues can be used to augment the differentiated function of MSC, and to provide a suitable extracellular environment to promote bone repair and integration, while also allowing guided and minimally-invasive delivery to defect sites.

## Supplementary Figures and Legends

**Supplementary Figure. 1.**
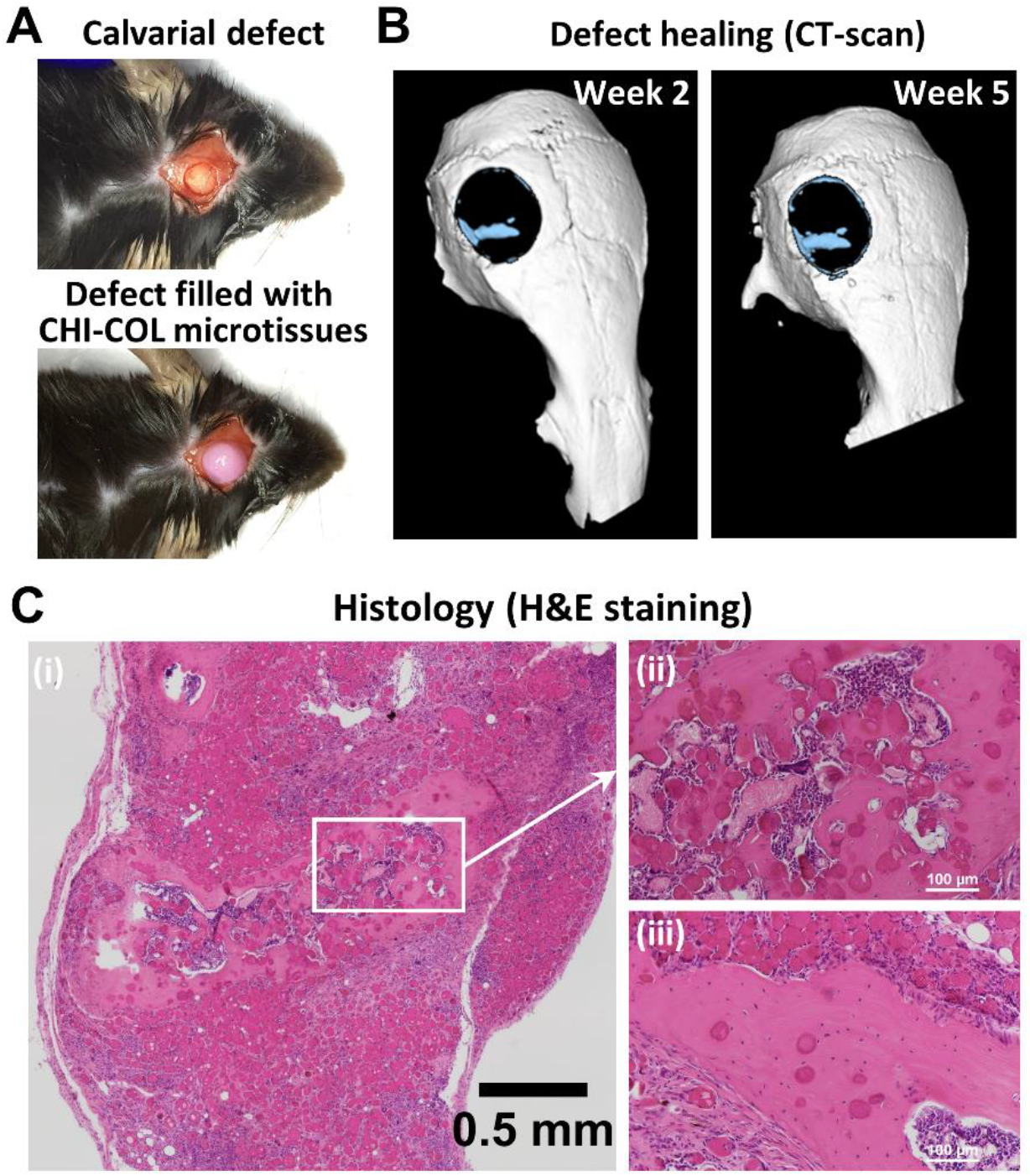
Screening of chitosan-collagen microtissue formulation. (A) Chitosan-collagen (CHI-COL) hydrogel formulation that can create stable microtissues was screened. The composite made up of 0.2 % collagen, and 0.25 % chitosan exhibited minimal contraction after the addition of glyoxal and β-glycerophosphate (B) Early peak in the expression of osteocalcin by day 14 in 0.25% chitosan composite indicated an increase in osteogenicity of microtissues with an increase in chitosan concentration. Based on our screening 0.2% collagen-0.25% collagen hydrogel composite was deemed suitable for fabricating osteogenic microtissues through emulsification. (C) The inclusion of HA particles increased the size of the microtissues. The acellular microtissues (0% HA) had a diameter of 54.3±12 μm while the diameter of 0.5%, 1% and 2% HA CHI-COL microtissues were 73.9±17.7 μm, 79.7±18.7 μm, and 148.6±21.4 μm respectively.

**Supplementary Figure 2.**
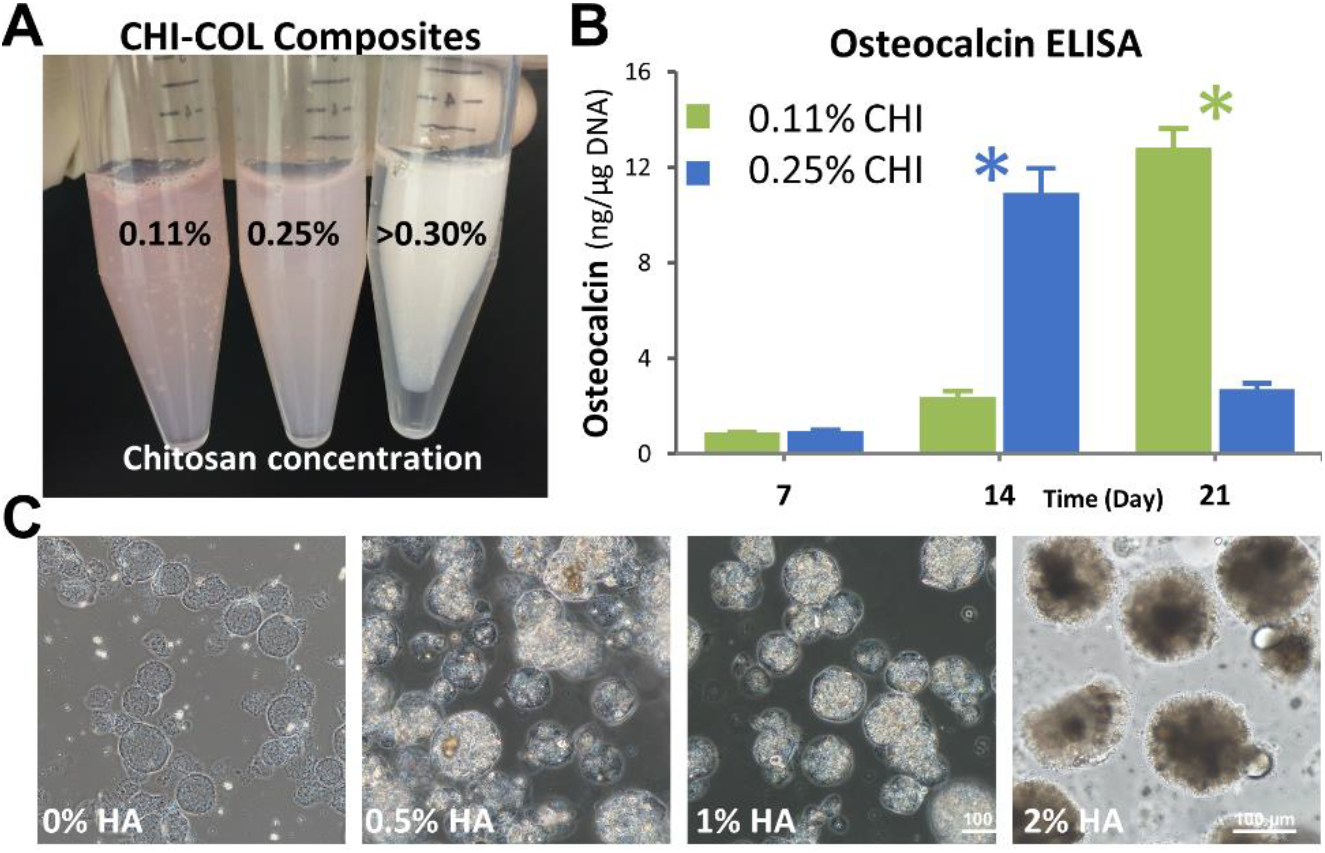
Temporal analysis of microtissues chemical structure using NMR analysis. The chemical composition of the microtissues seeded with MSC stayed intact and showed the presence of both collagen and chitosan for up to 21 days under culture conditions.

**Supplementary Figure 3.**
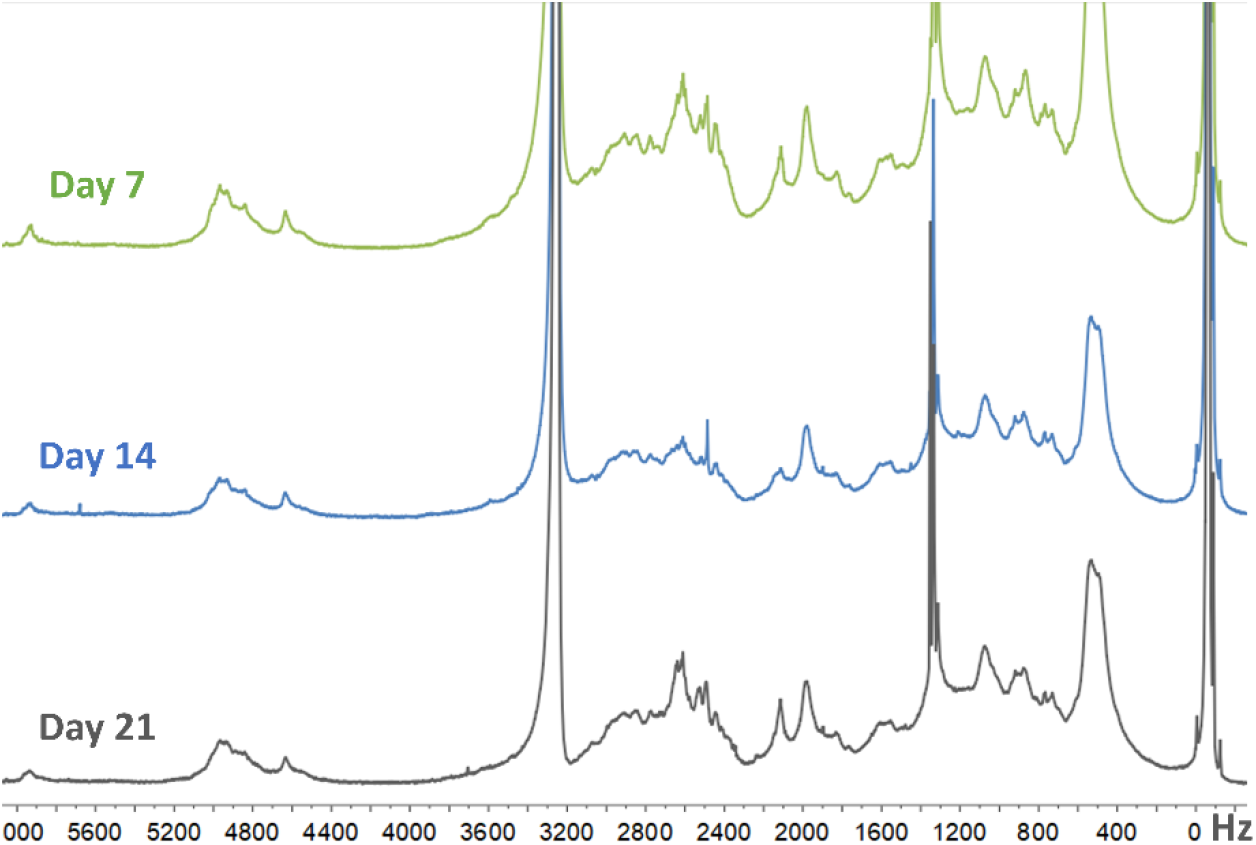
In-vivo characterization of COL-CHI microtissues without HA. (A) A unilateral 4 mm full-thickness critical-sized calvarial defect was created using a diamond-coated trephine bit in the non-suture associated right parietal bone, taking care to avoid dural injury. The implant was squirted to fill the defect, and the skin was sutured immediately. (B) The microCT scan of the defect at weeks 2 and 5 showed very low ossification and wound closure. (C) Histological analysis of calvarial samples at week-6. (i) The H&E staining of the sections revealed areas of the lamellar bone formation with remnants of the microtissues embedded within the structure (ii) and (iii). However, CHI-COL microtissues with MSC couldn’t completely heal the defect with calcified bone.

**Supplementary Figure 4.**
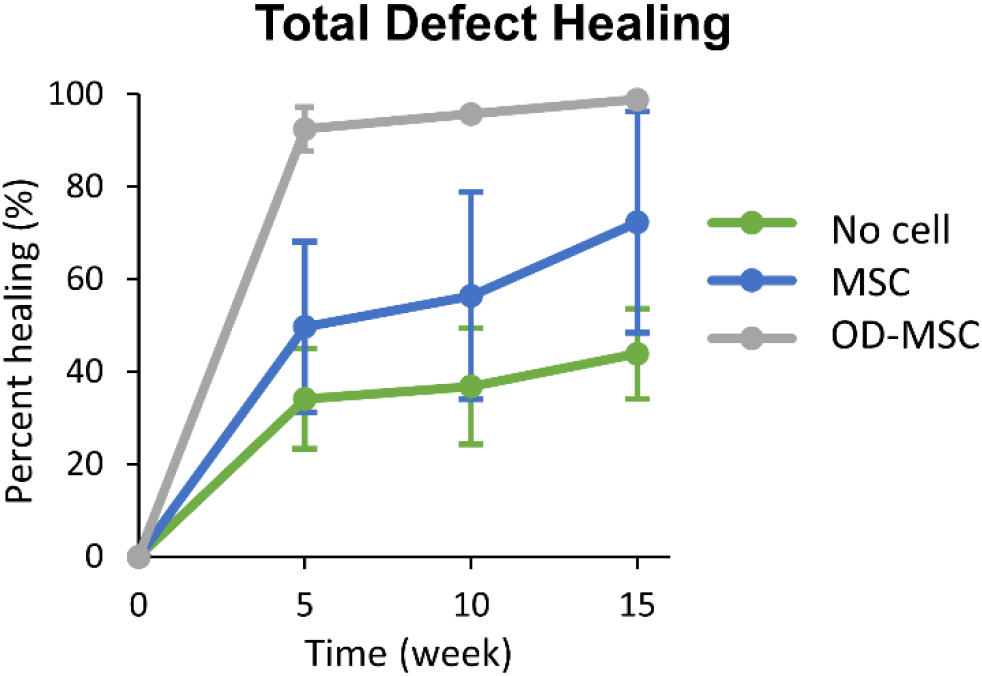
Quantification of defect healing through image analysis. Gross examination of the 3D reconstructed μCT images showed that by week 10, the OD-MSC microtissues had bridged >95% of the defect area. In contrast, microtissues with MSC showed only ~50% coverage, and microtissues without cells showed the lowest bridging of the defect measuring only ~30% of the defect area.

**Supplementary Figure 5.**
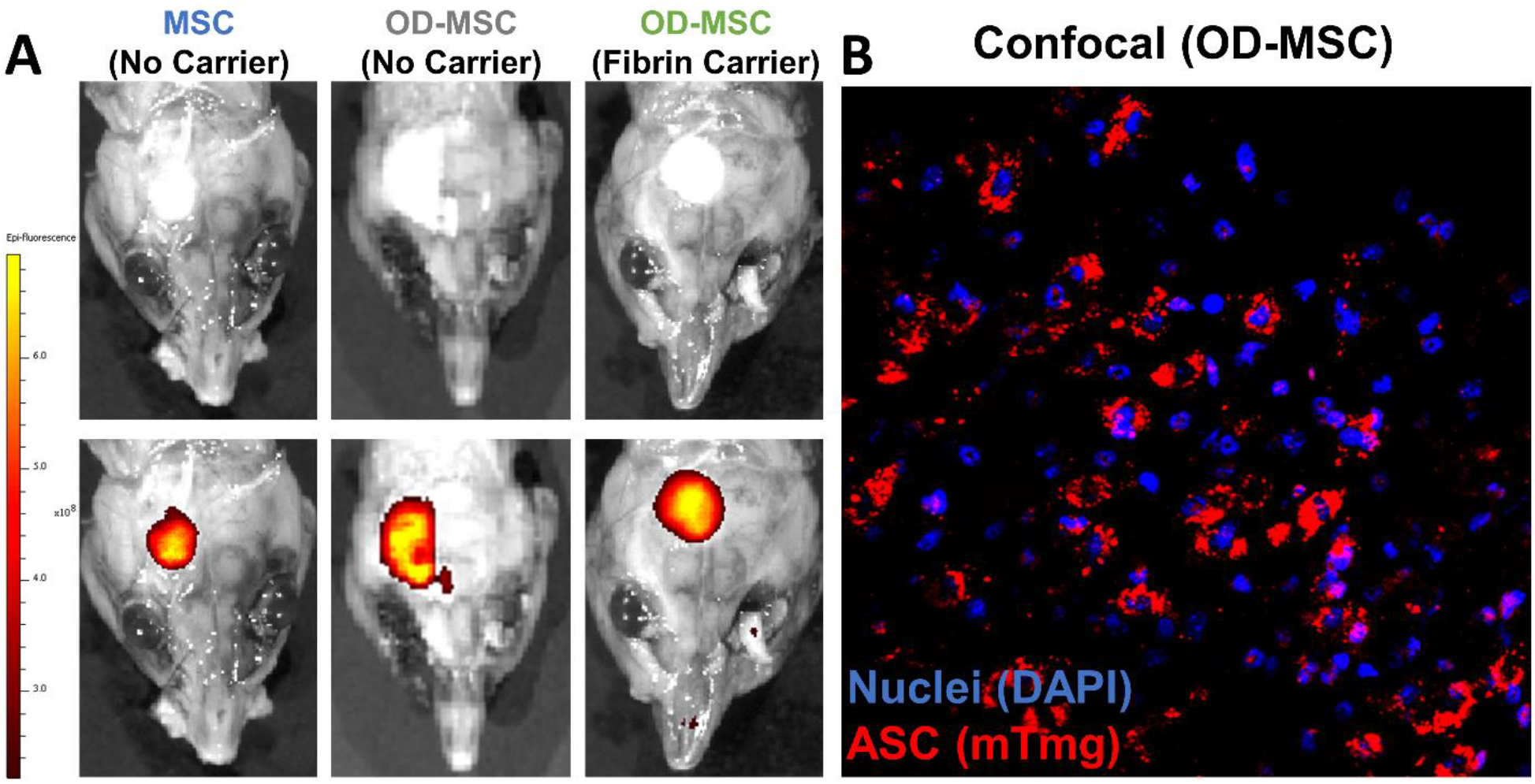
Fluorescent imaging of the implanted MSC within the constructs at week 12. The IVIS^®^ Spectrum in vivo imaging of the calvaria shows the presence of the red-fluorescent exogenous mTmG MSC in all conditions, confirming the engraftment of the supplied MSC. The intensity of the fluorescence shows the concentration of cells in the defect area. (B) Confocal fluorescent image of the OD-MSC construct with fibrin carrier gel show the presence of red-fluorescent mTmG MSC. The cells were counterstained with DAPI nuclear stain.

**Supplementary Figure 6.**
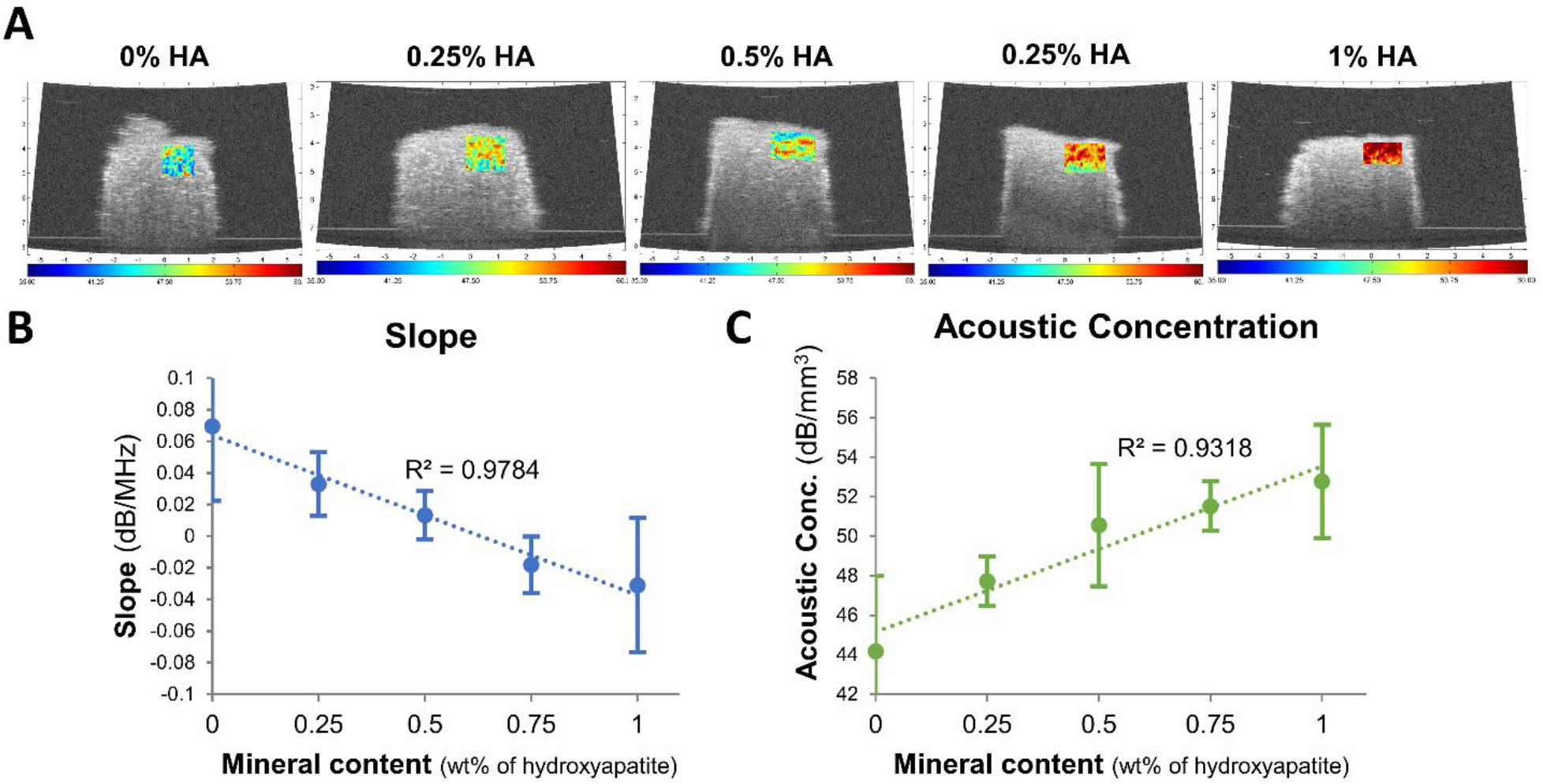
The spectral ultrasound imaging (SUSI) to quantify the mineral content of the implanted tissues. (A) The Grayscale images of the constructs made up of microtissue with 0.0 to 1.0% HA overlaid with the corresponding acoustic concentration values show an upward trend. (B) A plot of the mineral content of the microtissues versus the slope of the plot showed substantial agreement (R=0.97). (C) A plot of the mineral content of the microtissues versus the acoustic concentration showed substantial agreement (R=0.93).

